# How do your genetics compare to the world’s elite marathoners?

**DOI:** 10.1101/369066

**Authors:** Gregory E. Sims, Xiang-pei Zeng, Jeffrey Falk, Michael Nova

## Abstract

Pathway OME (Formerly Pathway Genomics, http://www.pathway.com) has identified a unique set of genetic traits associated with endurance, anxiety and likelihood of hamstring/Achilles injuries in an elite set of marathon runners. To date, this is the largest database of elite marathon athletes to ever have their exomes sequenced. Proprietary SportIQ Artificial Intelligence (AI) algorithms were developed and used to compare genetic traits in these 1,119 elite marathon runners to a normal population enabling us to develop a ‘Marathon Runner’ profile to assist athletes in training to compete and achieve their athletic goals. Athlete’s whose genetic profiles differ significantly from the “Marathon Runner” profile will gain additional insights on the specific training necessary to prepare their muscles for the demanding conditions of long distance running.

## Introduction

The performance characteristics of elite endurance athletes such as marathoners are believed to have a strong genetic basis (Tucker *et al* 2013). The genetic predisposition is believed to comprise a significant portion of athletic performance in ‘power’ athletes, with greater than 60%, of performance tied to genetics and only one third due to environmental components such as training, nutrition and technological aids (Weyerstraß *et al*, 2018). The prototypical distance runner tends to be shorter in height, lighter in weight, with less muscular calves and limited muscle mass in the upper body. These physical characteristics embody an economical human form able to sustain high speeds for continuous time periods. Several well studied genes have a measurable impact on physical capacity, such as genes known to contribute to physical endurance, maximal aerobic capacity (VO_2_ max), and lactic acid clearing after exercise (Sarzynski et 2017) A runner with the right combinations of essential genes, might exhibit an innate talent for running that provides advantages over other athletes with less optimal genetics. Furthermore, examining a runner’s genetic profile can provide insights into whether he or she should focus training and goals towards sprints, mid-distance or marathons.

Pathway OME has developed the SportIQ genetic test which focuses on collecting actionable insights on sports health and performance based on your genetics. A portion of the test profiles your genetic similarity to world class marathoners. This marathon profile was built using data that Pathway OME collected at an elite qualification-based international marathon held in the United States in 2015. We collected exome genotyping data across 230,000 genetic variations for 1,119 mixed-race runners who prequalified for the marathon. Genetics results of these elite marathoners were analyzed with a subset of the SportIQ AI algorithm alongside a set of 2,500 normal individuals to determine what distinguishes these runners from a normal population group (mixed race).

## Methods

We Exome genotyped 1,119 individuals who pre-qualified (elite-status) for an international marathon in the United States using the Illumina’s Infinium HumanCoreExome-12v1.1 Chip. The microarray chips were analyzed using the iScan System. The resulting genotyping data was analyzed using a custom designed AI-driven algorithm, SportsIQ-ex which uses a 13-marker subset of the markers which compose the SportIQ Test. The SportsIQ-ex Algorithm predicts phenotypic outcome data in seven categories: Endurance, Power, Achilles Tendinopathy, Recovery Time, Hamstring Injury, Bone Density and Calcium Uptake, and Anxiety, using genetic risk factors from published, human GWAS studies. SportIQ-ex was developed to collect sports performance insights using only the exonic markers present on the 12v1.1 Chip. We compared the marathoner population to the publicly available genotyping data for 2,500 HAPMAP cell lines. This collection of HAPMAP individuals serves as a normal control population. Using the relative differences in genotype frequencies between the two datasets of selected markers from the SportIQ-ex algorithm we developed a weighted score model to report the genetic similarity between an individual’s genetic profile and an optimal Marathoner. We describe the method of calculating the Marathon score here: Let *m_i_* = { *aa*, *ab*, *bb*} represent the frequency values for the three possible zygosity states (aa=homozygous reference, ab=heterozygous, bb=homozygous alteration) for marker *i* as observed in the Marathon individuals. Likewise let *h_i_* represent the genotype frequencies for marker *i* in the HAPMAP set. We use the frequency difference between *m*_i_ and *h_i_* to derive a score, *s_i_*, for each observable genotype, where 
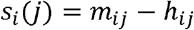
and the index *j* represents one of the 3 possible genotypes. Now, for each query sample we wish to score, we calculate the sum of scores for *N* markers given the specific genotype present in the query *q*:

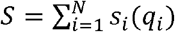

The effect of this scoring is to weight genotypes more frequently observed in the Marathon dataset. The individual score contributions of a specific genotype can range from positive to negative in value. For example, rs699 which contributes to the ‘Power’ phenotype, has a score contribution of 0.13 for the A/A genotype and −0.27 for the G/G genotype. We used this score function to rescore all HAPMAP samples to determine if there are any ethnicity related trends. This score function is also forms the basis of our Marathon Score phenotype in our commercially available SportIQ test.

## Results

### Phenotype outcome comparison

We compared the results of the marathon phenotypes and genotypes to the normal controls. The phenotype outcome comparisons (Figure 1) show significant differences in a number of phenotypes: Endurance (*p*<0.01), Achilles injury risk (*p*<0.01) and anxiety (*p*<0.01).

**Figure 1:**
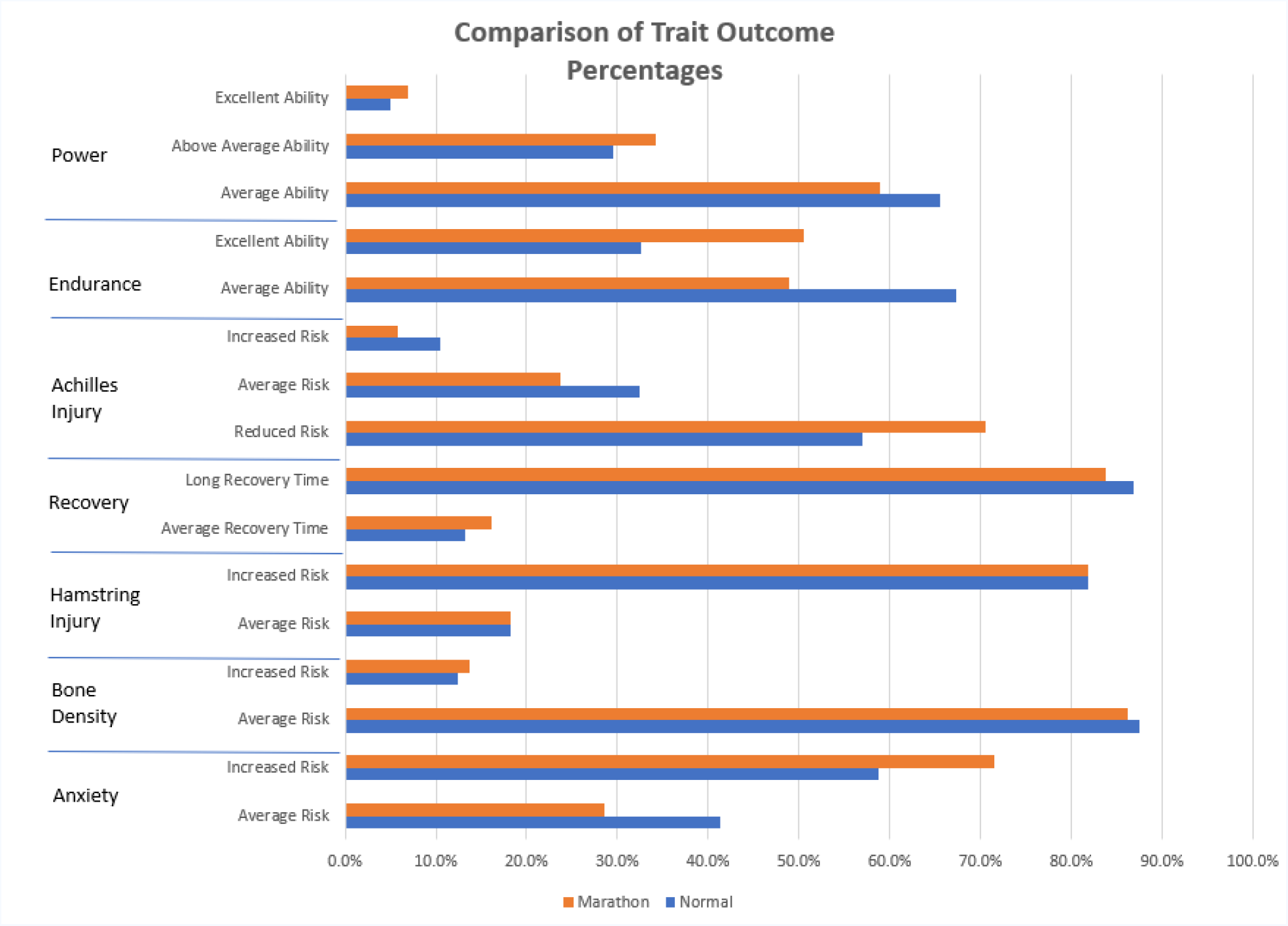
Distribution of individual Phenotype/trait outcomes from the Sports-ex algorithm. The percentage of the population of the 1,119 marathoners (orange) and 2,500 normal individuals (blue) is shown for several SportsIQ phenotypes. Note significantly different outcomes for marathoners in endurance, Achilles injury and anxiety.

### Sports Endurance

In general we predict that marathoner population will have a higher percentage of individuals with excellent endurance ability. We found some surprising results with relation to known endurance markers. Specifically, ADRB2 rs1042713 (allele A) was previously found to be associated with faster marathon times (Tsainos et al, 2010). In our dataset marathoners tended to have a higher proportion of individuals without this variation. Likewise, AMPD1 rs17602729 (allele G), also has been implicated in increased endurance (Ginevičienė et al 2014). We found that a higher proportion of marathoners in our dataset were only carriers of this allele (G;A genotypes).

### Sports Power

Intuitively, marathoners wouldn’t be expected to have genetics associated with ‘power’ athlete status. Weight lifters and rowers need quick bursts of power, from non-propulsive muscle; for the marathoner that extra muscle is just an additional non-economical burden. Our comparisons show this to be true to some extent. Marathon runners are similar in power marker distribution to the normal population. A few markers in the marathon population are notable because they tend to show power disinclination. For example, individuals who carry homozygote G allele (G;G) of the SOD2 rs4880 are associated with power athlete status (Weyerstraß *et al*, 2018), while we found that marathon runners have tend to not carry the G allele.

### Achilles Tendinopathy

The Achilles tendon is a band of tissue that connects the calf muscle to the heel bone. Achilles tendinopathy is characterized by inflammation and lesions in the tendon. Our study shows that marathoners have a strongly reduced risk of developing this type of injury due to genetic conditions. Individuals who were homozygous for the C allele of the rs679620 in *MMP3* had 2.5-times higher risk of Achilles tendinopathy than other individuals (Raleigh *et al*, 2009). Marathoners tend to not be carriers of the C allele. Achilles **tendinitis** tends to affect runners more than any other group or athletic population (Sheehan 1997), so it naturally follows that long distance runners would have some natural reduced risk, which our study shows.

### Recovery Time

During exercise, contracting skeletal muscles produce lactate and hydrogen ions as a result of glycolysis. If the individual has slower lactate removal from muscles, the recovery time may be longer after exercise or training. Individuals carrying the A allele of rs1049434 in the *MCT1* gene are associated with slower lactate clearance during muscle recovery and therefore longer recovery time (Fedotovskaya *et al*, 2014). Marathoners have no significantly different expected recovery time than the normal population.

### Hamstring Injury

The hamstring muscle is the large muscle that pulls on the hamstring tendon. This tendon attaches to the large muscles at the back of the thigh to bone. The main mechanisms of hamstring injury is an extreme contraction of the muscle at high velocity or slow stretching at the outermost range of motion (Malliaropoulos et al, 2018). Several of the genes associated with Achilles Tendon injuries are also associated with hamstring injuries. Individuals carrying the *TNC* risk allele T (A allele reported in the paper) at rs2104772 are associated with reduced risk of Hamstring injury (Larruskain *et al*, 2018). Marathoners show no significantly greater risk of hamstring injuries.

### Bone Density and Calcium Uptake

Osteoporosis is a metabolic bone disease, characterized by low bone mass and bone tissue deterioration, which leads to osteoporotic fracture risk. Individuals who carrying homozygote A allele (A;A) rs2228570 in *VDR* are associated with lower bone mineral density (Yasovanthi *et al*, 2011). Individuals who receive an outcome of “Increased Risk” are recommended to increase their calcium (CA) intake. The risk of Osteoporosis and low bone density in marathon runners is no different than the normal population.

### Anxiety

An interesting observation of this study is that we found that marathoners are predicted to have a significantly higher risk of anxiety. Specifically, individuals carrying the risk allele A at the rs4680 locus of *COMT* gene are associated with risk of anxiety (Enoch *et al*, 2008). A significantly higher proportion of marathon runners are homozygous (A;A) for rs4680. A;A often called the ‘worrier’ (as opposed to ‘warrior’) genotype characterized by lower pain thresholds, enhanced vulnerability to stress and may be associated with advantages in tasks requiring focus. It is tempting to speculate that there is a connection between focus/determination and the willpower need to complete a marathon event.

### Identifying strong potential Marathoners in HAPMAP

We employed our marathon profiling algorithm to identify key groups of HAPMAP individuals who might have the genetics to be marathoners. Figure 2 show the distribution of scores across all HAPMAP individuals used in this study. Table I provides more details about the highest individuals in the uppermost scoring bin from Figure 2. The ethnic breakdown is surprising. In recent Olympic games, especially the Rio de Janeiro games of 2016 – the distance races are heavily dominated by athletes from East African nations. However no East African haplogroups are in the top bin. We examined the samples collected for Luhya (HAMAP code: LWK) population of Kenya and found this population has a mean score of −0.70 +/- 0.05. We compared this to the British group (GBR), Utah residents (CEU) and Han Chinese (CHS) groups and found respective scores of 0.03 +/- 0.05, 0.02 +/- 0.05, -.61 +/- 0.05. The ethnic breakdown of the Marathon runner dataset is unknown and would be useful for further stratification. Also, there are some inherent limitations in the studies we used to construct our SportIQ algorithm since much of the literature evidence is derived from studies on limited population sets, with specific genetic variations only studied in Caucasians, Asians and African Americans.

**Figure 2:**
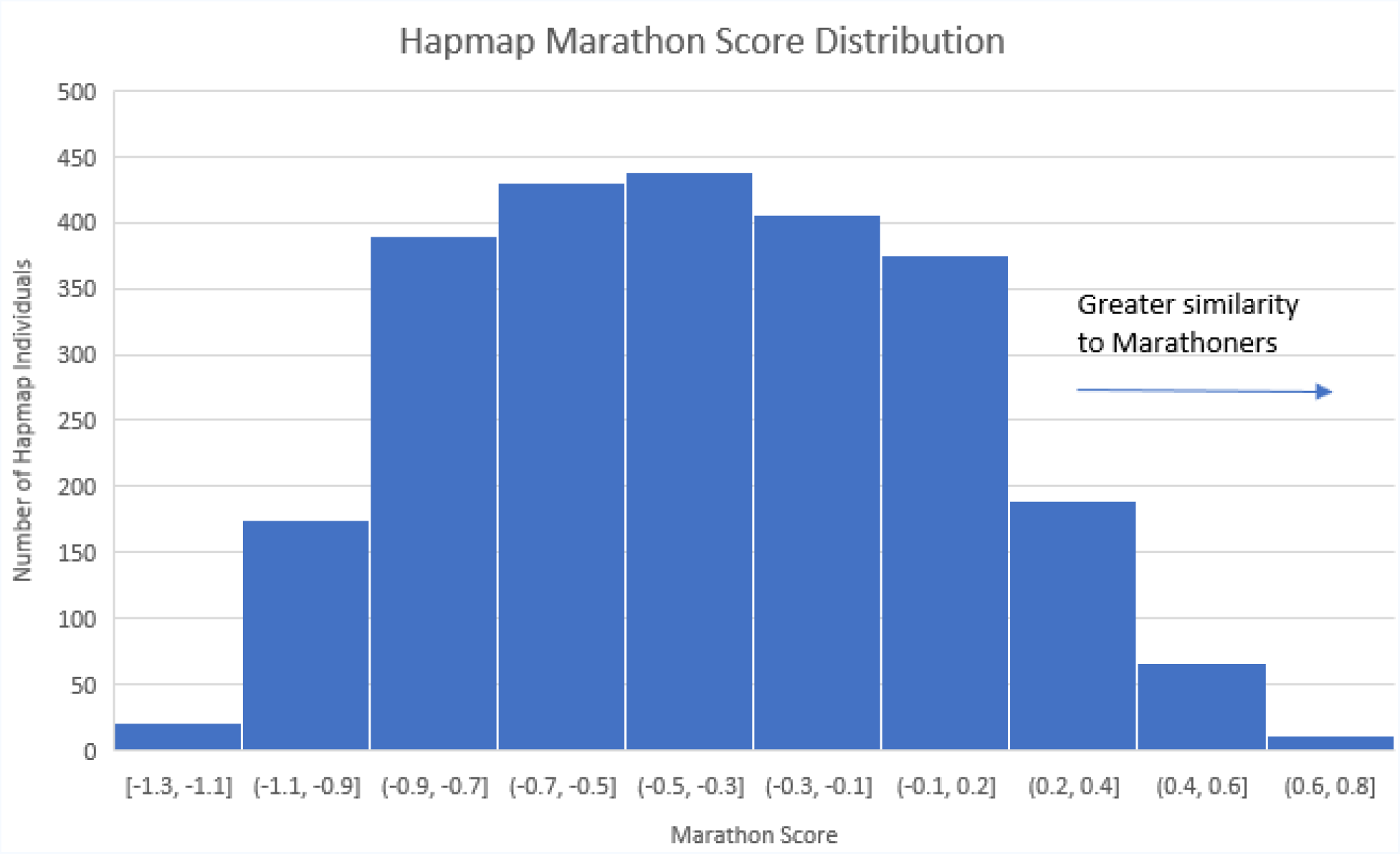
Distribution of Hapmap Individuals by Marthon Score. Table I represents a selection of the hapmap individuals scoring in the 0.6 to 0.8 score interval.

**Table I.** Profile of top scoring HAPMAP genotypes.

| Table I. Profile of top scoring HAPMAP genotypes |  |  |  |  |  |
| --- | --- | --- | --- | --- | --- |
| HAPMAP ID | Code | Population | Superpopulation | Sex | Marathon Score |
| HG00264 | GBR | British | European | Male | 0.79 |
| NA06985 | CEU | CEPH | European | Female | 0.77 |
| NA12890 | CEU | CEPH | European | Female | 0.72 |
| NA20803 | TSI | Tuscan | European | Male | 0.70 |
| HG00183 | FIN | Finnish | European | Male | 0.69 |
| NA21128 | GIH | Gujarati | South Asian | Male | 0.66 |
| HG01623 | IBS | Spanish | European | Female | 0.66 |
| HG01670 | IBS | Spanish | European | Female | 0.65 |
| HG01168 | PUR | Puerto Rican | American | Female | 0.65 |
| NA20772 | TSI | Tuscan | European | Female | 0.63 |
| HG04222 | ITU | Indian | South Asian | Male | 0.62 |

## Conclusion

Our marathon score profile was built using data that Pathway OME collected at an elite qualification-based international marathon event and represents what we believe to be the largest exome-dataset of marathon runners collected to date. We see significant differences in this marathon population that are related to genetic traits associated with endurance, anxiety and likelihood of hamstring/Achilles injuries. We have made these genetic insights available to the public through our SportsIQ test which we anticipate will be available July of 2018. Further in-depth comparison analysis of individual genetic markers of the marathon runners verses normal mixed-race population databases will be demonstrated in follow-on publications.

